# A new phylogenetic protocol: Dealing with model misspecification and confirmation bias in molecular phylogenetics

**DOI:** 10.1101/400648

**Authors:** Lars S Jermiin, Renee A Catullo, Barbara R Holland

## Abstract

Molecular phylogenetics plays a key role in comparative genomics and has an increasingly-significant impacts on science, industry, government, public health, and society. In this opinion paper, we posit that the current phylogenetic protocol is missing two critical steps, and that their absence allows model misspecification and confirmation bias to unduly influence our phylogenetic estimates. Based on the potential offered by well-established but under-used procedures, such as assessment of phylogenetic assumptions and tests of goodness-of-fit, we introduce a new phylogenetic protocol that will reduce confirmation bias and increase the accuracy of phylogenetic estimates.

**Dedication:** To the memory of Rossiter H. Crozier (1943-2009), an evolutionary biologist, who, with his great generosity and wide-reaching inquisitiveness, inspired students and scientists in Australia, and abroad.

## Introduction

Molecular phylogenetics plays a pivotal role in the analysis of genomic data and has already had a significant, wide-reaching impact in science, industry, government, public health, and society (Table 1). Although the science and methodology behind applied phylogenetics is increasingly well understood within parts of the scientific community (45), there is still a worryingly large body of research where the phylogenetic analysis was done with little attention to the consequences of a statistical misfit between the phylogenetic data and the assumptions that underpin the phylogenetic methods.

**Table 1.** Examples of phylogenetic research, divided into areas, based on impact and/or relevance.

| Area | Examples |
| --- | --- |
| Science | Provide accurate estimates of the evolution of, for example, species (1-4)<br>Provide accurate estimates of evolutionary processes at the molecular level (5,6)<br>Addressing macroevolutionary questions pertaining to birds and mammals (7,8)<br>Understand biogeographic patterns and diversity (9,10)<br>Reconstruction of ancestral states (11,12) |
| Industry | Facilitate the design and engineering of novel enzymes (13) and drugs (14-16) |
| Government | Reveal likely sources and dispersal routes of agricultural pests and pathogens (17-21)<br>Assign conservation priorities to species or biogeographic regions based on estimates of genetic diversity (22-24) |
| Public health | Reveal the origin and spread of human pathogens (25-28)<br>Predict the evolution of human influenza A (29)<br>Identify the natural reservoir of the virus causing COVID-19 (42-44)<br>Reveal the origin and evolution of cancers (30,31) |
| Society | Reveal the evolution of human language (32,33)<br>Map the relationship among ancient texts (34), tales (35), and music (36)<br>Reveal evolution of humans since their divergence from other primates (37-41) |

One reason for this is that phylogenetics relies extensively on statistics, mathematics, and computer science, and many users of phylogenetic methods find the relevant sections of these disciplines challenging to comprehend. Another reason is that methods and software often are chosen because they already are popular or easy to use, rather than because they are the most appropriate for the scientific questions and phylogenetic data at hand. A third reason is that much of the phylogenetic research done to date has relied on phylogenetic protocols (46–51), which have evolved to become a standard to which it seems sensible to adhere. Although these protocols vary, they have, at their core, a common set of sensible features (i.e., procedural steps) that are linked in a seemingly logical manner (see below).

Here we posit that, although the current phylogenetic protocol has many useful features, it is missing two crucial components whereby the quality of fit between the data and models applied is assessed. This means that using the phylogenetic protocol in its current form may lead to biased conclusions. We suggest a modification to the protocol that will make it more robust and reliable, and that will identify where new methods are needed to enable ease of use and statistical accuracy in the field.

### The current phylogenetic protocol

Phylogenetic analysis of alignments of nucleotides or amino acids usually follows a protocol like that in Figure 1. Initially, the phylogenetic data are chosen on the assumption that they will allow the researchers to solve a particular scientific problem. This choice of sequences data is often based on prior knowledge, developed locally or extracted from the literature. Then, a multiple sequence alignment (MSA) method is chosen, often on the basis of prior experience with a specific method. The sequences are then aligned, the aim being to obtain an MSA, wherein homologous characters (i.e., nucleotides or amino acids) are aligned. In practice, it is often necessary to insert alignment gaps between some of the characters in some of the sequences to obtain an optimal MSA—in some cases, there may be characters in different sequences that cannot be aligned reliably.

**Figure 1.**
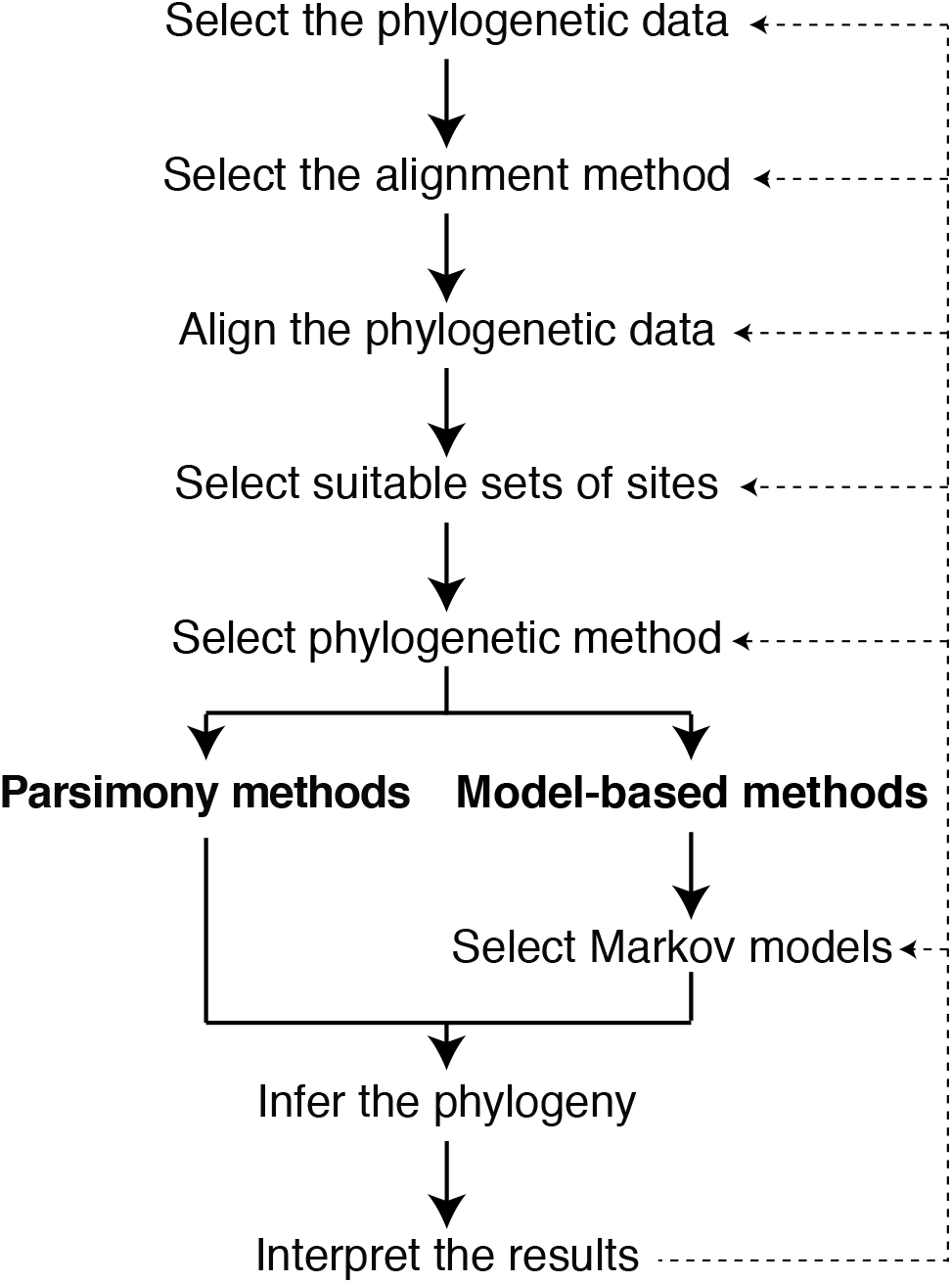
The current phylogenetic protocol. Solid arrows show the order of actions normally taken during a phylogenetic analysis. Dashed arrows show feedback loops often employed in phylogenetic research. For details, see the main text.

Then follows the task of selecting sites that will be used to infer the phylogenetic tree. The rationale behind doing so is to maximize the signal-to-noise ratio in the MSA. By omitting poorly-aligned and highly-variable sections of the MSA, which are thought to create noise due to the difficulty of establishing true homology for each site (defined as similarity due to historical relationships by descent (52)), it is hoped that the resulting sub-MSA will retain a strong historical signal (defined as the order and timing of divergence events (53)) that will allow users to obtain an accurate phylogeny. The choice of sites to retain is made by visual inspection of the MSA or by using purpose-built software (54–64). The automated ways of filtering MSAs have been questioned (65).

Having obtained a sub-MSA, the next step in the protocol is to select a phylogenetic method. The choice of phylogenetic method implies accepting the assumptions on which the method rests. For example, irrespective of whether the data comprise nucleotides, codons, or amino acids, it is often assumed that the sequences evolved along a single bifurcating tree and that the evolutionary processes operating at the variable sites in the sequences are independent and distributed identically (in a probabilistic sense). If model-based molecular phylogenetic methods are chosen, the underlying assumption usually is that the evolutionary processes operating at the variable sites can be approximated accurately by using Markov models that are stationary, reversible, and homogeneous (66–68) over time (the assumption of evolution under SRH conditions) (for more details about the most common phylogenetic assumptions,see Box 1). In practice, the choice is one between methods assuming that the underlying evolutionary processes can be modelled using a Markov model of nucleotide or amino-acid substitutions (i.e., distance methods (69–74), likelihood methods (69,71,73–79), Bayesian methods (80–87)), or non-parametric phylogenetic methods (i.e., parsimony methods (69,71,73,74,88–90)). In reality, many researchers analyze their data using a range of modelbased phylogenetic methods, and reports that only use parsimony methods are increasingly rare. Depending on the chosen phylogenetic method, researchers may have to select a suitable model of sequence evolution (i.e., a model that combines the substitution model and the rates-across-sites model) to apply to the sub-MSA. This choice is often made by using model-selection methods (6,91–102).

Having chosen a phylogenetic method and, in relevant cases, a suitable model of sequence evolution, the next step involves obtaining accurate estimates of the tree and evolutionary processes that led to the data. Phylogenetic methods are implemented in many software packages (69–85,88–90) and, depending on the methods chosen, users often also obtain the nonparametric bootstrap probability (103) or clade credibility (104) to measure support for divergence events in the phylogeny.

Having inferred the phylogeny, the final step in the protocol is to interpret the result. Under some conditions—most commonly, the inclusion of out-group sequences—the tree may be drawn and interpreted as a rooted phylogeny, in which case the order of divergence events and the lengths of the individual edges may be used to infer, for example, tempo and mode of evolution of the data. Often, the inferred phylogeny confirms earlier-reported or assumed evolutionary relationships. Often, too, there are surprises, which are difficult to understand and explain. If the phylogenetic estimate is convincing and newsworthy, the discoveries may be reported, for example, through papers in peer-reviewed journals.

On the other hand, if the surprises are too numerous or unbelievable, the researchers may begin the task of finding out what may have ‘gone wrong’ during the phylogenetic analysis.

This process is depicted as dashed feedback loops in Figure 1. The researchers may analyze their data differently (e.g., use other Markov models, use other phylogenetic methods, use a different sub-MSA, align the sequences differently, use a different alignment method, or use another data set), and given enough patience, they may reach a conclusion about their data and attempt to publish their results.

### Problems with the current phylogenetic protocol

Although the current phylogenetic protocol has led to many important discoveries, it also has left many scientists with strong doubts about or, alternatively, undue confidence in the estimates. The literature is rife with examples where analyses of the same data have led to disagreements among experts about what is the ‘right’ phylogeny (cf. e.g., (105–107)). Such disagreements are confusing, especially for non-experts and the public. To understand why these disagreements might arise, it is necessary to understand the challenges that applied phylogenetic research still faces.

While it is clear that the right data are needed to answer the scientific question at hand, making that choice is not always as trivial as it might seem. In some cases, the sequences may have evolved too slowly and/or be too short, in which case there may not be enough information in the data, or they have evolved so fast that the historical signal has largely been lost (108). In rarely-reported cases, the data are not what they purport to be (109).

Next, there is no consensus on what constitutes an optimal MSA. In simple terms, what is required is an accurate MSA where every site is a correct homology statement. Currently, there is no automatic procedure for assessing homology (52). Frequently, different methods return different MSAs, implying different homology statements, but they cannot all be right. Averaging MSAs has been proposed as a solution (110). Producing an MSA will often involve manual modifications following visual inspection, which introduces subjectivity and a lack of reproducibility. One way to mitigate this problem is to rely on reviews and simulation-based comparisons of MSA methods (52,111–118), but these reports seem to have had less impact than deserved. It is clear that poor MSAs can cause problems for downstream analyses such as inference of positive selection (119), ancestral state reconstruction (120), and estimates of phylogeny (121).

Having identified an MSA, the choice of poorly-aligned and/or highly-variable sites to omit (in a process commonly referred to as masking (64)) depends not only on the MSA method used but also on how difficult it is to detect these sites—it is impractical to visually inspect MSAs with more than ~50 sequences and ~300 sites. In the past, expert knowledge about the data was often used (e.g., structural information about the gene or gene product), but automated methods (54–64) are now frequently used. However, these methods often yield different sub-MSAs from the same MSA, leaving confusion and doubt.

The choice of what phylogenetic method to use for the data is rated (by many) as the most challenging one to make (e.g., because the assumptions underpinning each phylogenetic method often are poorly understood), and it is often solved by using several phylogenetic methods. If these methods return the same phylogenetic tree, many authors feel confident that they have identified the ‘true’ phylogeny and they would go on to publish their results. However, while this approach may have led to correct trees, it is perhaps more due to luck than to scientific rigor that the right tree was identified. This is because every phylogenetic method is based on assumptions (see above), and if these assumptions are not violated too strongly by the data, and the number of variable sites is sufficient, then the true tree has a high probability of being identified. On the other hand, if the violations are strong enough, there is currently no way of knowing whether the correct tree has been identified. Indeed, strong violation of phylogenetic assumptions could lead to similar, but nevertheless, wrong trees being inferred using different phylogenetic methods (122,123).

Over the last two decades, the choice of a suitable model of sequence evolution has often been made by using purpose-built model-selection methods (6,91–102). Assuming a tree, these methods step through a list of predefined models, evaluating each of them, one by one, until the list is exhausted. This approach is sensible if the true or most appropriate model is included in the list of predefined models. On the other hand, if the true model is not included in this list, then the popular model-selection methods will never be able to return the true model. They will return an optimal model, but it will be conditional on the models included in that list. Unfortunately, most model-selection methods only consider time-reversible Markov models. If the sequences have evolved along a single tree but under non-reversible Markovian conditions (i.e., under non-SRH conditions), then there is no way that a single, time-reversible Markov model is sufficient to approximate the evolutionary processes across all edges of the tree (124). Therefore, it is worrying that many researchers still ignore or dismiss the implication of compositional heterogeneity across sequences (123). This type of heterogeneity indicates that the evolutionary processes has changed across the lineages. This implication must be taken seriously when data are analyzed phylogenetically.

The choice of phylogenetic program is often driven by prior experiences and transaction costs (i.e., the time it takes to become a confident and competent user of the software) rather than by a profound understanding of the strengths, limitations, and weaknesses of the available software. However, this may not substantially minimize the accuracy of the phylogenetic estimate, as long as the data do not violate the assumptions on which the phylogenetic methods are based, and the phylogenetic methods search tree space and model space thoroughly.

The bootstrap probability (103) and clade credibility (104) are often thought of as metrics of the accuracy of the phylogenetic estimate or the confidence we might have in the inferred divergence events. Unfortunately, doing so is unwise because they measure consistency of the estimate (125)—a phylogenetic estimate may consistently point to an incorrect tree.

Finally, once well-supported phylogenetic estimates have been inferred, prior expectations are likely to influence whether the estimates are considered both reliable and newsworthy. In some cases, where information on the phylogeny is known (e.g., serially-sampled viral genomes), not meeting the prior expectations may signal a problem with the phylogenetic analysis. However, if a researcher’s expectations are met by the phylogenetic results, it is more likely that a report will be written without a further assessment of what might have gone wrong during the analysis. This tendency—allowing prior expectations to influence the interpretation of phylogenetic estimates—is called confirmation bias. Confirmation bias is not discussed in phylogenetics, even though it is a recognized problem in other disciplines (e.g., psychology and social science (126)), so it is timely that the phylogenetic community takes onboard the serious implications of this.

### The new phylogenetic protocol

Although the current phylogenetic protocol has many shortcomings, it also has many good attributes, including that it is easy to apply and implement as a pipeline. But to mitigate its limitations, it will be necessary to redesign the protocol to accommodate well-established, but largely-ignored, procedures as well as new feedback loops.

Figure 2 shows a picture of the new phylogenetic protocol. It shares many features found in the current protocol (e.g., the first four steps). However, the fifth step (assess phylogenetic assumptions) will be novel to many researchers. As all phylogenetic methods are based on assumptions, it is sensible to validate these assumptions at this point in the protocol. Since many phylogenetic methods assume that the data (e.g., different genes) have evolved over the same tree, and that the chosen data partitions have evolved independently under the same time-reversible Markovian conditions, it is wise to survey the sub-MSAs for evidence that the sequences actually have evolved under these conditions. If the data violate these phylogenetic assumptions, then it will be wise to avoid these phylogenetic methods and to employ other such methods. Alternatively, it may be worth following the relevant feedback loops in Figure 2—perhaps something led to a biased sub-MSA? The relevance and benefits of this step are illustrated using a case study (Box 2), which focuses on determining whether a data set is consistent with the phylogenetic assumption of evolution under time-reversible conditions. Assessments of other phylogenetic assumptions require other types of tests and surveys. Some of these relevant questions and methods are listed in Table 2 (53,127–150).

**Figure 2.**
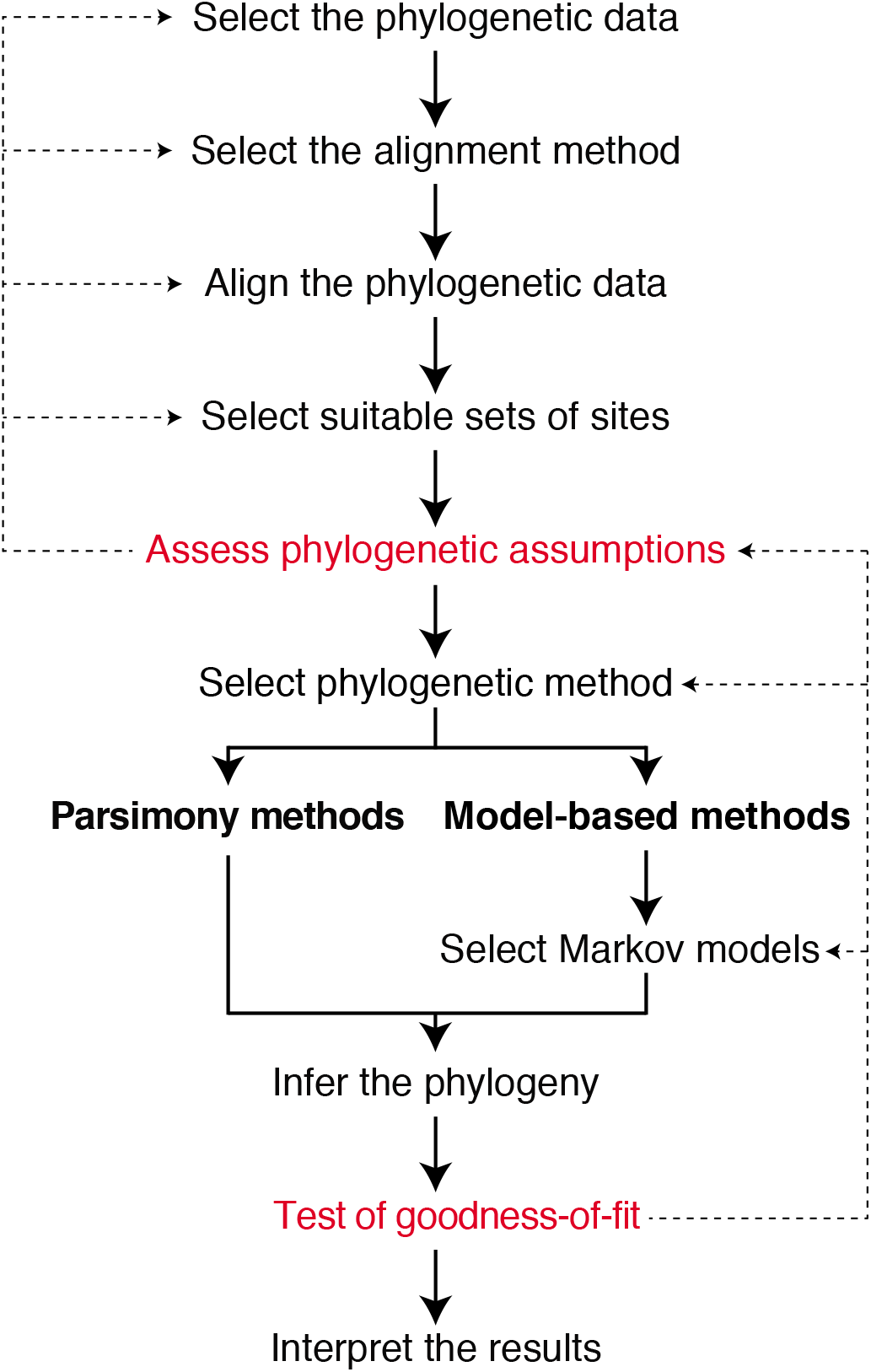
A new phylogenetic protocol. Solid arrows show the order of actions normally taken during a phylogenetic analysis. Dashed arrows show feedback loops often employed in phylogenetic research. For details, see the main text.

**Table 2.**
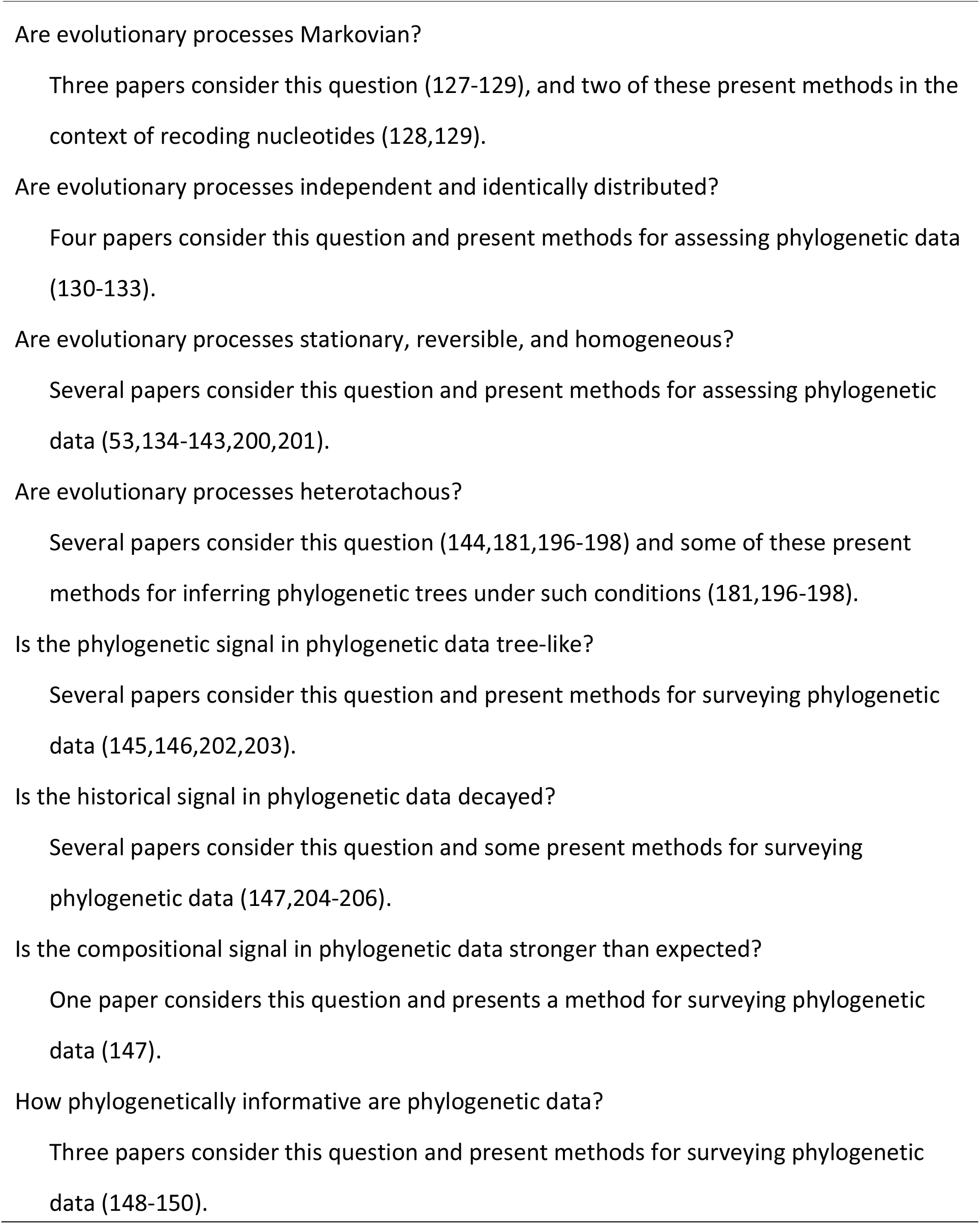
Important questions requiring pre-phylogenetic surveys of phylogenetic data, accompanied by publications describing methods developed to carry such surveys

Next follows the choice of phylogenetic method, but now this choice is made on the basis of the previous step, rather than on cultural or computational reasons. If the sequences have evolved on a single tree under time-reversible Markovian conditions, there is a large set of phylogenetic methods to choose from (69–85,88–90). On the other hand, if these data have evolved under more complex Markovian conditions, the number of suitable phylogenetic methods is, frustratingly, rather limited (5,67,151–176), and most of these methods are aimed at finding the optimal model of sequence evolution for a given tree rather than finding the optimal set of trees. Users of phylogenetic methods therefore are sometimes confronted by a dilemma: Do they abandon their data set because it has evolved under nontime-reversible conditions and because there are no appropriate phylogenetic methods for such data, or do they take the risk and employ the phylogenetic methods that assume evolution under time-reversible conditions? Fortunately, there may be a way around this dilemma.

Having inferred the phylogeny using model-based phylogenetic methods, it is possible to test the fit between tree, model and data (step 10 of the new protocol). A suitable test of goodness-of fit was proposed in 1993 (177) (Fig. 4). In brief, using the inferred optimal tree, including the edge lengths, it is possible to simulate data sets under the null model (i.e., the inferred optimal model of sequence evolution with its parameter values included). This is called a parametric bootstrap. Given the optimal tree and the optimal model of sequence evolution, several sequence-generating programs (5,165,178–181) facilitate production of pseudo-data. Having generated, say, *m* = 1,000 pseudo-data, the next step involves finding the difference (δ) between the unconstrained log-likelihood (i.e., the log-likelihood inferred without assuming a tree and a model of sequence evolution) and constrained log-likelihood (i.e., the log-likelihood inferred assuming a tree and a model of sequence evolution)—that is, computing δ = *lnL*(**D**) — *lnL*(**D**|*T, M*), where **D** is the data, *T* is the optimal tree, and *M* isthe optimal model of sequence evolution. If δ is greater for the real data than it is for the pseudo-data, then that result reveals a poor fit between tree, model, and data (141). The approach described here works well for likelihood-based phylogenetic analysis. A similar approach that relies on posterior predictive distributions is available for Bayesian-based phylogenetic analysis (182).

**Figure 3.**
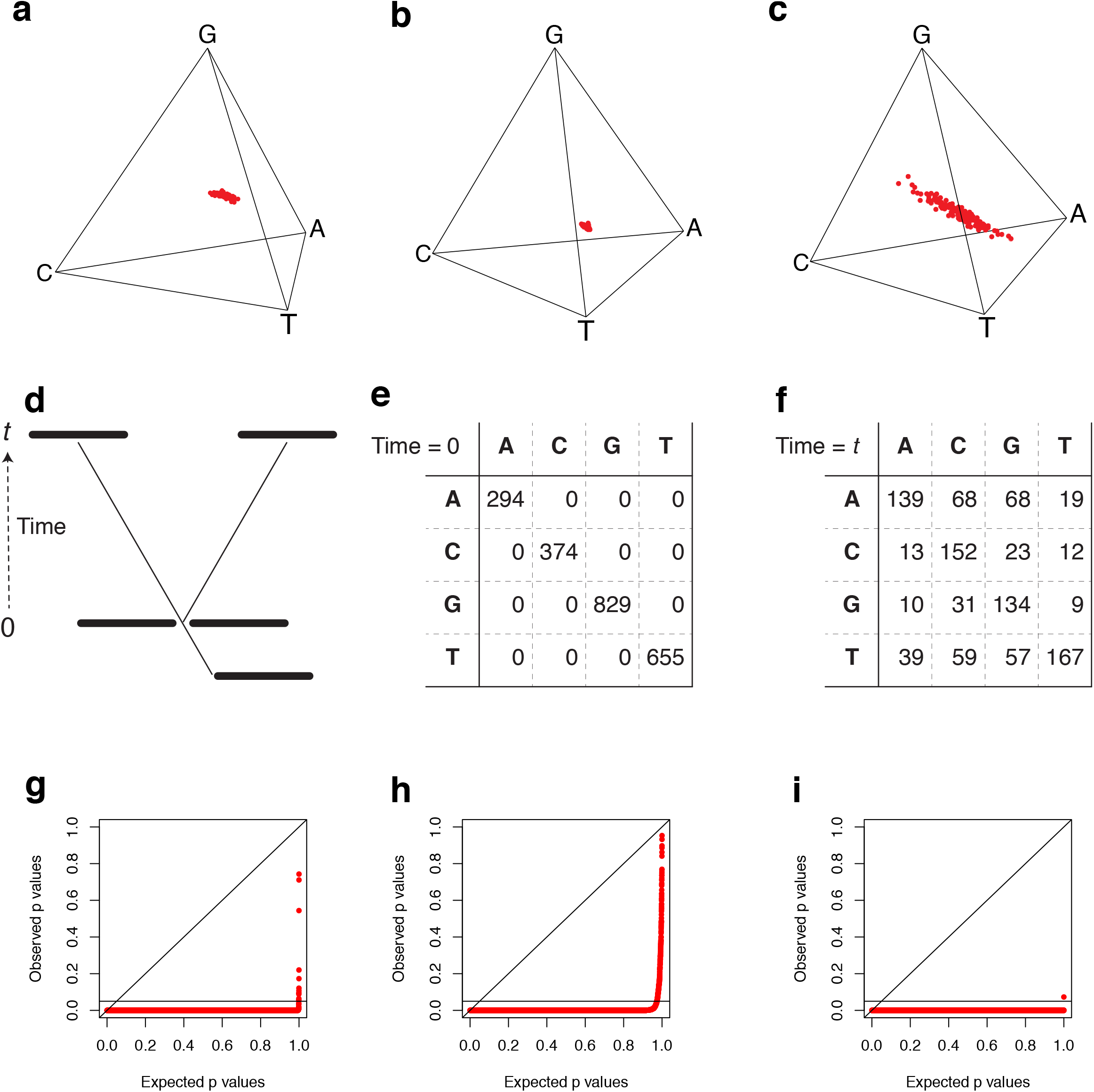
Illustrating the merits of the fifth step in the new phylogenetic protocol. Panels **a-c** show the average nucleotide composition at first, second, and third codon position of the sequences (144 sequences, 413,459 codons) originally examined by Misof et al. (3). Each dot represents the nucleotide composition of a single sequence. The spread of dots in panels **a-c** reveals compositional heterogeneity at first, second, and third codon position, indicating these data violate important assumptions underlying most phylogenetic methods. The tetrahedral plots were generated using SeqVis (139). Panel **d** shows a two-tipped tree with an ancestral sequence and two diverged sequences. Panels **e** and **f** show the divergence matrix for these sequences at time 0 and time *t*. Each number in a cell of a divergence matrix corresponds to the number of sites with nucleotide *i* in one sequence, and nucleotide *j* in the other. Panels **h-i** shows the PP plots for the data already analysed in panels **a-c**. A total of 10,296 tests were done for each of the three codon positions using Homo 1.3 (http://www.csiro.au/homo/ — a new version of Homo is now available (53)).

**Figure 4.**
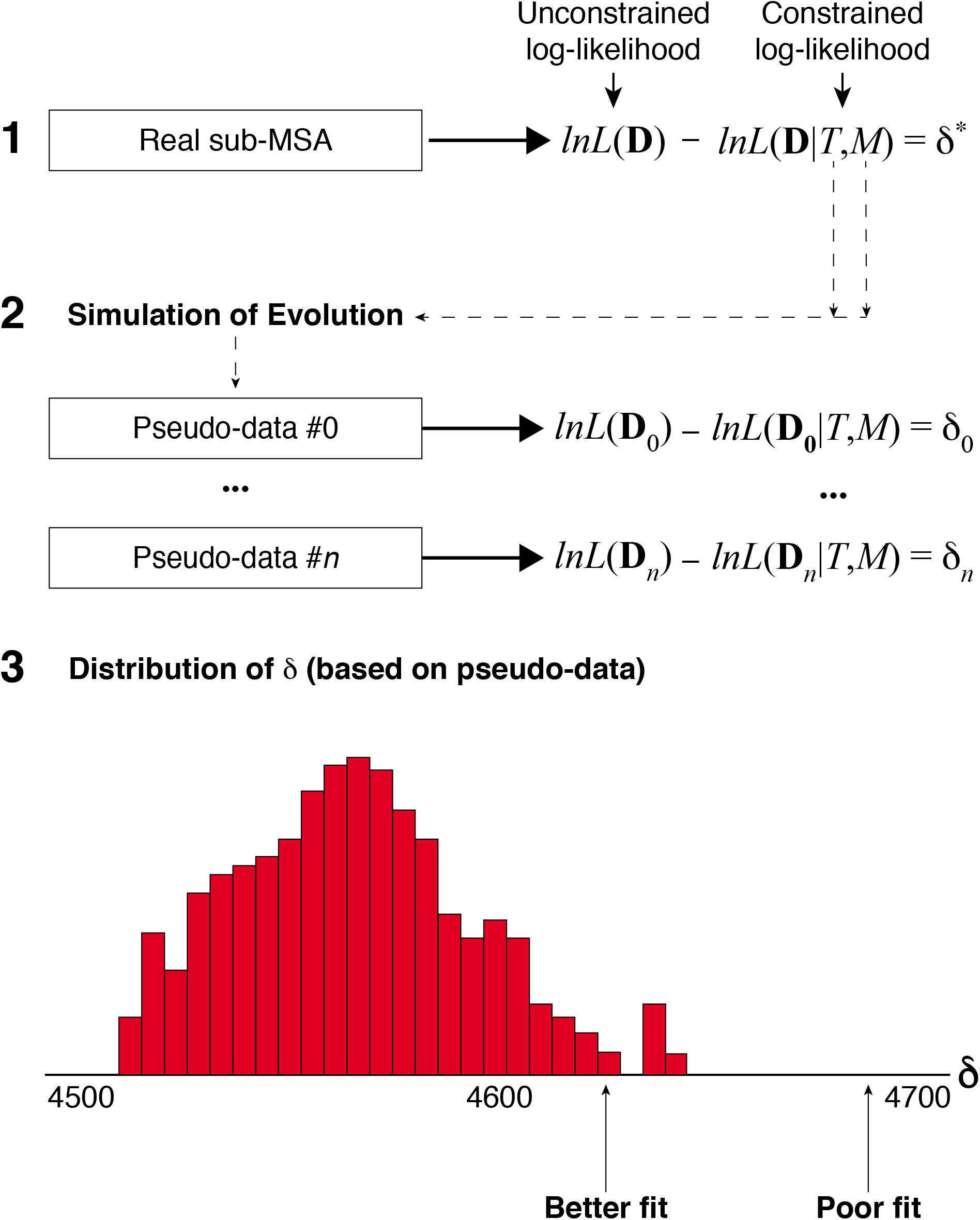
Diagram showing the parametric bootstrap procedure that may be used to conduct a suitable goodness-of-fit test. The procedure includes three steps. For details, see the main text.

Parametric bootstrapping is computationally expensive and may be time-consuming, so we recommend it be used only if the data appears to meet the assumptions of the phylogenetic method used. The advantages of using such a goodness-of-fit test is that it allows users to assess whether a poor fit is poor enough to not be due to chance. It does not say anything about why a fit might be poor or whether the lack of fit matters (in a biological sense). If the fit is poor, then the relevant feedback loops should be followed (Fig. 2)—perhaps a biasing factor was overlooked? If the phylogenetic tree and model of sequence evolution are found to fit the data, then that implies that the estimates represent a plausible explanation of the data. It is these estimates that should be reported, but only as one plausible explanation, not as the only possible explanation. This is because there may be other plausible explanations of the data that never were considered during the analysis.

### The future: Areas in most need of methodological research

Adherence to the new phylogenetic protocol will undoubtedly lead to improved accuracy of phylogenetic estimates and a reduction of confirmation bias. The advantage of the fifth step in the new phylogenetic protocol (i.e., assess phylogenetic assumptions) is that users are able to decide how to do the most computationally-intensive parts of the phylogenetic study without wasting precious time on, for example, a high-performance computer centre. Model selection, phylogenetic analysis, and parametric bootstrapping are computationally-intensive and time-consuming, and there is a need for new, computationally efficient strategies that can be used to analyse sequences that have evolved under complex phylogenetic conditions.

The advantage of the tenth step in the new phylogenetic protocol (i.e., test goodness-of-fit) is its ability to answer whether an inferred phylogeny explains the data well, or not. In so doing, this step tackles the issue of confirmation bias front on. Clearly, without information gleaned from the fifth step, the parametric bootstrap might return an unwanted answer (i.e., the inferred tree and model of sequence evolution does not fit the data well), so to avoid this disappointment it is better to embrace the new phylogenetic protocol in full.

Results emerging from studies that rely on the new phylogenetic protocol might well call into question published phylogenetic research, but there is also a chance that research might gain stronger support. This is good for everyone concerned, especially since it will become easier to defend the notion that the research was done without prejudice or preference for a particular result. Objectivity should be restored in phylogenetics—it is no longer reasonable to defend phylogenetic results on the basis that they were obtained using the best available tools; if these tools do not model the evolutionary processes accurately, then that should be reported rather than be hidden away. This is critical because it increases the transparency of the research and aids other scientists to understand the intricate nature of the challenges encountered.

Notwithstanding the likely benefits offered by the new protocol and the methods supporting it, it would be unwise to assume that further development of phylogenetic methods will no longer be needed. On the contrary, there is a lot of evidence that method development will be needed in different areas:

1. *MSA Methods—There* is a dire need for MSA methods that yield accurate homology statements. Likewise, there is a need for methods that allow users to: (*i*) determine how accurate different MSA methods are, and (*ii*) select MSA methods that are most suitable for the data at hand. Moreover, there is a dire need for better transparency in the way MSAs are reported in the literature (64).
2. *Methods for Masking MSAs*—Assuming an accurate MSA has been inferred, there is a need for strategies that can be applied to identify and distinguish between poorly-aligned and highly-variable regions of MSA. Well aligned but highly-variable regions of MSAs may be more informative than poorly-aligned regions of such MSAs, so to delete them may be unwise.
3. *Model-selection Methods—Model* selection is a pivotal prerequisite if parametric phylogenetic methods are used. However, the model-selection methods currently employed may not be accurate (183), especially for sequences that have evolved under complex conditions (e.g., heterotachous, covarion, or non-time-reversible conditions). For example, the evolutionary process may have to be considered an evolving entity in its own right. Further, a better understanding of the information criteria used is necessary (184).
4. *Phylogenetic Methods—While* there are many accurate phylogenetic methods for analysis of data that have evolved under time-homogeneous, reversible Markovian conditions, there is a dearth of accurate phylogenetic methods suitable for analysis of data that have evolved under more complex conditions. Added to this challenge are methods that accurately consider incomplete lineage sorting of genetic markers and the special conditions associated with the analysis of SNP data.
5. *Goodness-of-fit Tests*—Suitable goodness-of-fit tests are available, but there is not only a need for a wider understanding of the merits of these tests, but also of how they can be tailored to suit different requirements. In particular, there is a need for programs that can generate simulated data under extremely complex evolutionary conditions. Some programs are available (5,165,180,185), but they only cater for a very limited set of conditions.
6. *Analysis of Residuals—Although* goodness-of-fit tests can tell you whether or not the lack of fit observed is potentially due to chance, they do not answer the more useful question of whether or not that lack of fit matters or how the lack of fit arises (186,187). For this reason, residual diagnostic tools that can inform the user about the way in which their model fails to fit the data would be very useful.

In summary, while calls for better phylogenetic methods and more careful considerations of the data have occurred (124), we posit there is a need for a comprehensive overhaul of the current phylogenetic protocol. The proposed new phylogenetic protocol is unlikely to be a final product; rather, it is probably a first, but important step towards a more scientifically sound phylogenetic protocol, which not only should result in more accurate phylogenetic estimates and but also to a reduction in the likelihood of confirmation bias.

## Conclusions

The Holy Grail in molecular phylogenetics is being able to obtain accurate, reproducible, transparent, and trustworthy phylogenetic estimates from the phylogenetic data. We are not there yet, but encouraging progress is being made in not only in the design of the phylogenetic protocol but also in phylogenetic methodology based on the likelihood and Bayesian optimality criteria.

Notwithstanding this progress, a quantum shift in attitudes and habits is needed within the phylogenetic community—it is no longer enough to infer an optimal phylogenetic estimate. The fit between trees, models, and data must be evaluated before phylogenetic estimates can be considered newsworthy. We owe it to the scientific community and wider public to be as rigorous as we can—the attitude “She’ll be alright, mate” is no longer appropriate in this discipline.

## ACKNOWLEDGEMENTS

L.S.J. thanks the University College Dublin for its generous hospitality. We thank D. Higgins, A. Locatelli, and K. H. Wolfe for their constructive feedback.

## AUTHOR CONTRIBUTIONS

L.S.J. conceived the new phylogenetic protocol. R.A.C., B.R.H., and L.S.J. wrote the paper.

## COMPETING FINANCIAL INTERESTS

The authors declare no competing financial interests.

### BOX 1 COMMON PHYLOGENETIC ASSUMPTIONS

Phylogenetic methods developed to analyze alignments of nucleotides, codons, and amino acids typically make some of all of the following modelling assumptions:

1. The sequences evolved along the edges of a single bifurcating tree
2. Each site has a fixed rate of mutation that does not change across the tree; however, note that this rate can be zero (invariable sites),
3. The evolutionary processes operating at the variable sites follow independent and identically-distributed Markov processes, and
4. The evolutionary processes operating at the variable sites were stationary, reversible, and homogeneous over time.

Other modelling assumptions are often made (e.g., that rate-heterogeneity across sites can be modelled accurately using a discrete Γ distribution (188) or nonparametric, probabilistic model of rate-heterogeneity across sites (189); the latter strategy is more flexible, allowing multimodal distributions of rate-heterogeneity across sites to be accommodated (6)), but they are not considered here. In addition to these assumptions, it is typically assumed that the data are phylogenetically informative—that is, the historical signal has not decayed enough to make it pointless using the data in a phylogenetic study. We now describe the modelling assumptions with a minimum of mathematical and statistical terminology.

#### The assumption of tree-like evolution—Building

on graph theory, this assumption—the treelikeness assumption—states that every site in an alignment has evolved on the same rooted phylogenetic tree. The assumption applies equally to single-gene and multi-gene data sets. Some relatively-recent methods (190–192) relax this assumption by assuming a single species tree but allowing the gene trees to vary from the species tree due to incomplete lineage sorting. However, if, for example, some of the sequences in the data were involved in recombination events or introgression, then the phylogenies of the sites on either side of the break points will be different and the assumption of a single underlying tree will be violated.

#### The assumption of each site having a fixed rate of mutation that does not change across the tree

Building on knowledge of biochemistry and natural selection, the sites in a nucleotide sequence can be divided into those that are variable and those that are invariable. Here the variable sites are those that are free to change while the invariable sites are those that cannot change, usually due to all mutations being deleterious.

A key point here is that it is not possible, in general, to distinguish between sites that were able to change but have not done so, and sites that were unable to change due to selection. For this reason, we are limited in our ability to conclude whether sites are truly invariable. Therefore, many phylogenetic methods are designed to estimate the proportion of invariable sites. In fact, it is highly recommended to assume that a proportion of sites are invariable.

The assumption that sites are either variable or invariable may be violated by the data if the selective constraints at the sites change over time. Under this scenario, a variable site might become invariable within, for example, one lineage but remain variable in all other lineages. Our understanding of the prevalent of evolving selective constraints is not good, but intra- and inter-molecular epistasis might be contributing factors. Intra- and inter-molecular epistasis is known to constrain the order and reversibility of amino acid substitution (193–195) but it has not yet been incorporated in phylogenetic methods (for exceptions, see 181,196,197,198).

#### The assumption of independent and identically-distributed Markovian processes

Based on probability theory, this assumption is required because it gives us the statistical power that is needed to compare evolutionary hypotheses within a statistical framework. A process is Markovian if the conditional probability of change at a site in a sequence depends only on the current state and is independent of its earlier states. In other words, a Markov process has no memory. The ‘independency of the Markovian processes’ refers to the processes at the variable sites being independent of one another, and the ‘distributions of the Markov processes being identical’ refers to the conditional probability of a change from state *i* to state *j* (e.g., from A to G) being the same irrespective of what variable site is considered. In other words, the assumption states that the variable sites evolve independently of one another and that the evolutionary processes are not only Markovian but also identical across the variable sites.

#### The assumption of stationary processes

A Markov process is stationary when the marginal probabilities of the states that the process might take remain the same irrespective of time. In mathematical terms, a Markov process is stationary when the following condition holds:

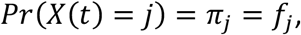

where *X*(*t*) denotes the state of the Markov process (*X*) at time *t, j* = A, C, G, T (for DNA), *π_j_* denotes the marginal probability of state *j* in the process, and *f_j_* represents the relative frequency of state *j* at the variable sites of the ancestral sequence. Usually, *π* is called the stationary distribution of the process (or the Markov model approximating this process). In mathematical terms, it is a vector.

The assumption of stationary processes is made as it reduces the number of parameters that needs to be optimised and, therefore, reduces the amount of time required to optimise them. If the sequences in an alignment have evolved under stationary conditions, then they are unlikely to have acquired different compositions of the states (e.g., nucleotides). On the other hand, if the sequences are compositionally heterogeneous, then they are unlikely to have evolved under stationary conditions. Evolution under non-stationary conditions now appears to be a norm rather than an exception (e.g., 142,199).

#### The assumption of reversible processes

A Markovian process is reversible if the probability of sampling state *i* from the stationary distribution (i.e., *π*) and going to state *j* is the same as the probability of sampling state *j* from the stationary distribution and going to state *j* In mathematical terms, a Markov process is reversible when the following condition holds:

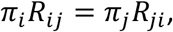

where *R_ij_* denotes the conditional probability of a substitution from state *i* to state *j*, and *i* = A, C, G, T (for DNA). In mathematical terms, *R* is a matrix. If the evolutionary process is consistent with this ‘balancing equation’, further reductions in the number of parameters can be made. An important upshot of this assumption is that the likelihood of a tree for a given phylogenetic data set can be computed without knowing the ancestral sequence for these data.

While the assumption of reversible processes is critical for most model-based phylogenetic methods, it is more difficult to ascertain whether sequences have evolved under reversible conditions. It is, however, clear that evolutionary processes cannot be reversible if they are non-stationary, so evolution under non-reversible conditions now also appears be a norm, rather than an exception. A biochemical study of the evolutionary trajectories between two enzymes, Atrazine chlorohydrolase (AtzA) and melamine deaminase (TriA), corroborates this picture: the optimal order of substitutions from AtzA to TriA differed from that from TriA to AtzA (195).

#### The assumption of homogeneous processes

A Markovian process is homogeneous in time if the conditional probabilities of substitutions are constant over time. When a Markov process is consistent with this assumption, it is easy to estimate the conditional probability of change (*P*) over a given time (Δ*t*). In mathematical terms, the following expression is used:

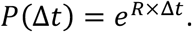

Here, *P* is a matrix representing the joint probability of state *i* at the beginning of an edge in a tree and state *j* at the end of this edge. By combining sets of *P* matrices, one per edge, in a manner that reflects the topology of the tree (i.e., *T*), it is possible to calculate the likelihood of the tree and the model (i.e., *R*), given the phylogenetic data.

Although convenient, the assumption of homogeneous processes over time is unlikely to be realistic. This is because accumulation of substitutions over time is likely to be gradual and occurring along the edges, rather than being punctuated and occurring only at the nodes in the tree. Sometimes, the term ‘homogeneous processes’ is used to describe evolutionary processes operating along diverging lineages in a phylogenetic tree. To avoid confusion, it is sensible to write ‘time-homogeneous processes’ when this is the intended meaning and to write ‘homogeneous processes across lineages’ when this is the intended meaning.

### BOX 2 CASE STUDY

To illustrate the relevance and benefits of the fifth step in the new phylogenetic protocol (i.e., assess phylogenetic assumptions), we surveyed the phylogenetic data used to infer the evolution of insects (3). The tetrahedral plots in Figure 3a–3c reveal that the nucleotide composition at the three codon positions is heterogeneous (most clear in panel c and least clear in panel b), implying that the evolutionary processes that operated at these positions are unlikely to have been time-reversible. However, the plots are deceptive because the presence of constant sites (i.e., sites with the same nucleotide or amino acid) in the data can mask how compositionally dissimilar the sequences actually are. To learn how to resolve this issue, it is necessary to focus on the evolution of two sequences on a tree (Fig. 3d) and the corresponding divergence matrix, **N**, at time 0 (Fig. 3e) and at time *t* (Fig. 3f). At time 0, the two sequences are beginning to diverge from one another, so the off-diagonal elements of **N** are all zero. Later, **N** may look like that in Figure 3f. The off-diagonal elements of **N** are now greater than zero, and matching off-diagonal elements of **N** might differ (i.e., *n_ij_* ≠ *n_ji_*). The degree of divergence between the two sequences can be inferred by comparing the off-diagonal elements to the diagonal elements, while the degree of difference between the two evolutionary processes can be inferred by comparing the above-diagonal elements of **N** to the below-diagonal elements of **N**. If the two evolutionary processes were the same, the matching off-diagonal elements in Figure 3f would be similar. A lack of symmetry (i.e., *n_ij_* ≠ *n_ji_*) implies that the evolutionary processes along the descendant lineages may be different. A matched-pairs test of symmetry (138) can be used to determine whether this observed deviation from symmetry is statistically significant. Figures 3g–3i show the distributions of the observed and expected *p* values from these tests for the data assessed in Figures 3a–3c. If the evolutionary processes operating along the descendant lineages are identical, the dots would be distributed along the diagonal line of a PP plot. However, as the dots in each plot fall far from this line, there is little evidence supporting the assumption of evolution under homogeneous conditions. The same is the case for the corresponding amino acid alignment (not shown), so it would be unwise to assume that the data evolved under time-reversible conditions. A more complex evolutionary process might explain these data. However, such methods were not available at the time when Misof et al. (3) obtained and analysed these data, and that is still the case!.

## Notes

### Competing Interest Statement

The authors have declared no competing interest.

## References

1. dos Reis, M., Inoue, J., Hasegawa, M., Asher, R.J., Donoghue, P.C.J. and Yang, Z.H. (2012) Phylogenomic datasets provide both precision and accuracy in estimating the timescale of placental mammal phylogeny. Proc. R. Soc. B, 279, 3491–3500.

2. Ruhfel, B.R., Gitzendanner, M.A., Soltis, P.S., Soltis, D.E. and Burleigh, J.G. (2014) From algae to angiosperms—inferring the phylogeny of green plants *(Viridiplantae)* from 360 plastid genomes. BMC Evol. Biol., 14, 26.

3. Misof, B., Liu, S.L., Meusemann, K., Peters, R.S., Donath, A., Mayer, C., Frandsen, P.B., Ware, J., Flouri, T., Beutel, R.G. et al. (2014) Phylogenomics resolves the timing and pattern of insect evolution. Science, 346, 763–767.

4. Prum, R.O., Berv, J.S., Dornburg, A., Field, D.J., Townsend, J.P., Lemmon, E.M. and Lemmon, A.R. (2015) A comprehensive phylogeny of birds (Aves) using targeted nextgeneration DNA sequencing. Nature, 526, 569–U247.

5. Jayaswal, V., Wong, T.K.F., Robinson, J., Poladian, L. and Jermiin, L.S. (2014) Mixture models of nucleotide sequence evolution that account for heterogeneity in the substitution process across sites and across lineages. Syst. Biol., 63, 726–742.

6. Kalyaanamoorthy, S., Minh, B.Q., Wong, T.F.K., Von Haeseler, A. and Jermiin, L.S. (2017) ModelFinder: fast model selection for accurate phylogenetic estimates. Nature Meth., 14, 587–589.

7. Penny, D. and Phillips, M.J. (2004) The rise of birds and mammals: are microevolutionary processes sufficient for macroevolution. Trends Ecol. Evol., 19, 516–522.

8. Meredith, R.W., Janečka, J.E., Gatesy, J., Ryder, O.A., Fisher, C.A., Teeling, E.C., Goodbla, A., Eizirik, E., Simão, T.L.L., Stadler, T. et al. (2011) Impacts of the Cretaceous terrestrial revolution and KPg extinction on mammal diversification. Science, 334, 521–524.

9. Knapp, M., Stockler, K., Havell, D., Delsuc, F., Sebastiani, F. and Lockhart, P.J. (2005) Relaxed molecular clock provides evidence for long-distance dispersal of Nothofagus (southern beech). PLoS Biol., 3, 38–43.

10. Jetz, W., Thomas, G.H., Joy, J.B., Hartmann, K. and Mooers, A.O. (2012) The global diversity of birds in space and time. Nature, 491, 444–448.

11. Marazzi, B., Ane, C., Simon, M.F., Delgado-Salinas, A., Luckow, M. and Sanderson, M.J. (2012) Locating evolutionary precursors on a phylogenetic tree. Evolution, 66, 3918–3930.

12. Pagel, M., Meade, A. and Barker, D. (2004) Bayesian estimation of ancestral character states on phylogenies. Syst. Biol., 53, 673–684.

13. Wilding, M., Peat, T.S., Kalyaanamoorthy, S., Newman, J., Scott, C. and Jermiin, L.S. (2017) Reverse engineering: transaminase biocatalyst development using ancestral sequence reconstruction. Green Chem., 19, 5375–5380.

14. Searls, D.B. (2003) Pharmacophylogenomics: Genes, evolution and drug targets. Nat. Rev. Drug Discov., 2, 613–623.

15. Goodfellow, M. and Fiedler, H.P. (2010) A guide to successful bioprospecting: informed by actinobacterial systematics. Antonie Van Leeuwenhoek, 98, 119–142.

16. Wright, G.D. and Poinar, H. (2012) Antibiotic resistance is ancient: implications for drug discovery. Trends Microbiol., 20, 157–159.

17. Boykin, L.M., Armstrong, K.F., Kubatko, L.S. and De Barro, P.J. (2012) Species delimitation and global biosecurity. Evol. Bioinform., 8, 1–37.

18. Hosokawa, T., Nikoh, N. and Fukatsu, T. (2014) Fine-Scale Geographical Origin of an Insect Pest Invading North America. PLoS One, 9, 5.

19. Yasaka, R., Ohba, K., Schwinghamer, M.W., Fletcher, J., Ochoa-Corona, F.M., Thomas, J.E., Ho, S.Y.W., Gibbs, A.J. and Ohshima, K. (2015) Phylodynamic evidence of the migration of turnip mosaic potyvirus from Europe to Australia and New Zealand. J. Gen. Virol., 96, 701–713.

20. Tay, W.T., Walsh, T.K., Downes, S., Anderson, C., Jermiin, L.S., Wong, T.K.F., Piper, M.C., Chang, E.S., Macedo, I.B., Czepak, C. et al. (2017) Mitochondrial DNA and trade data support multiple origins of Helicoverpa armigera (Lepidoptera, Noctuidae) in Brazil. Scientific Rep., 7, 45302.

21. Anderson, C.J., Oakeshott, J.G., Tay, W.T., Gordon, K.H.J., Zwick, A. and Walsh, T.K. (2018) Hybridization and gene flow in the mega-pest lineage of moth, Helicoverpa. Proc. Natl. Acad. Sci. USA, 115, 5034–5039.

22. Gonzalez-Orozco, C.E., Pollock, L.J., Thornhill, A.H., Mishler, B.D., Knerr, N., Laffan, S., Miller, J.T., Rosauer, D.F., Faith, D.P., Nipperess, D.A. et al. (2016) Phylogenetic approaches reveal biodiversity threats under climate change. Nat. Clim. Chang., 6, 1110–+.

23. Rosauer, D.F., Blom, M.P.K., Bourke, G., Catalano, S., Donnellan, S., Gillespie, G., Mulder, E., Oliver, P.M., Potter, S., Pratt, R. et al. (2016) Phylogeography, hotspots and conservation priorities: an example from the Top End of Australia. Biol Conserv, 204, 83–93.

24. Tucker, C.M., Cadotte, M.W., Carvalho, S.B., Davies, T.J., Ferrier, S., Fritz, S.A., Grenyer, R., Helmus, M.R., Jin, L.S., Mooers, A.O. et al. (2017) A guide to phylogenetic metrics for conservation, community ecology and macroecology. Biol. Rev., 92, 698–715.

25. Andersen, K.G., Shapiro, B.J., Matranga, C.B., Sealfon, R., Lin, A.E., Moses, L.M., Folarin, O.A., Goba, A., Odia, I., Ehiane, P.E. et al. (2015) Clinical sequencing uncovers origins and evolution of Lassa Virus. Cell, 162, 738–750.

26. Holmes, E.C., Dudas, G., Rambaut, A. and Andersen, K.G. (2016) The evolution of Ebola virus: Insights from the 2013-2016 epidemic. Nature, 538, 193–200.

27. Lanciotti, R.S., Lambert, A.J., Holodniy, M., Saavedra, S. and Signor, L.D.C. (2016) Phylogeny of Zika Virus in Western Hemisphere, 2015. Emerg. Infect. Dis., 22, 933–935.

28. Lessler, J., Chaisson, L.H., Kucirka, L.M., Bi, Q.F., Grantz, K., Salje, H., Carcelen, A.C., Ott, C.T., Sheffield, J.S., Ferguson, N.M. et al. (2016) Assessing the global threat from Zika virus. Science, 353, aaf8160.

29. Bush, R.M., Bender, C.A., Subbarao, K., Cox, N.J. and Fitch, W.M. (1999) Predicting the evolution of human influenza A. Science, 286, 1921–1925.

30. Alves, J.M., Prieto, T. and Posada, D. (2017) Multiregional tumor trees are not phylogenies. Trends Cancer, 3, 546–550.

31. Schwartz, R. and Schaffer, A.A. (2017) The evolution of tumour phylogenetics: principles and practice. Nat. Rev. Genet., 18, 213–229.

32. Pagel, M. (2009) Human language as a culturally transmitted replicator. Nat. Rev. Genet., 10, 405–415.

33. Bouckaert, R., Lemey, P., Dunn, M., Greenhill, S.J., Alekseyenko, A.V., Drummond, A.J., Gray, R.D., Suchard, M.A. and Atkinson, Q.D. (2012) Mapping the origins and expansion of the Indo-European language family. Science, 337, 957–960.

34. Barbrook, A.C., Howe, C.J., Blake, N. and Robinson, P. (1998) The phylogeny of The Canterbury Tales. Nature, 394, 839–839.

35. Tehrani, J.J. (2013) The phylogeny of Little Red Riding Hood. PLoS One, 8, 11.

36. Windram, H.F., Charlston, T. and Howe, C.J. (2014) A phylogenetic analysis of Orlando Gibbons’s Prelude in G. Early Music, 42, 515–+.

37. Ingman, M., Kaessmann, H., Pääbo, S. and Gyllensten, U. (2000) Mitochondrial genome variation and the origin of modern humans. Nature, 408, 708–713.

38. Ke, Y.H., Su, B., Song, X.F., Lu, D.R., Chen, L.F., Li, H.Y., Qi, C.J., Marzuki, S., Deka, R., Underhill, P. et al. (2001) African origin of modern humans in East Asia: A tale of 12,000 Y chromosomes. Science, 292, 1151–1153.

39. Schraiber, J.G. and Akey, J.M. (2015) Methods and models for unravelling human evolutionary history. Nat. Rev. Genet., 16, 727–740.

40. Posth, C., Renaud, G., Mittnik, A., Drucker, D.G., Rougier, H., Cupillard, C., Valentin, F., Thevenet, C., Furtwangler, A., Wissing, C. et al. (2016) Pleistocene mitochondrial genomes suggest a single major dispersal of Non-Africans and a late glacial population turnover in Europe. Curr. Biol., 26, 827–833.

41. Nielsen, R., Akey, J.M., Jakobsson, M., Pritchard, J.K., Tishkoff, S. and Willerslev, E. (2017) Tracing the peopling of the world through genomics. Nature, 541, 302–310.

42. Wang, H., Pipes, L. and Nielsen, R. (2020) Synonymous mutations and the molecular evolution of SARS-Cov-2 origins. BioRxiv, (https://doi.org/10.1101/2020.1104.1120.052019).

43. Boni, M.F., Lemey, P., Jiang, X., Lam, T.T.-Y., Perry, B., Castoe, T., Rambaut, A. and Robertson, D.L. (2020) Evolutionary origins of the SARS-CoV-2 sarbecovirus lineage responsible for the COVID-19 pandemic. BioRxiv, (https://doi.org/10.1101/2020.1103.1130.015008).

44. Liu, P., Jiang, J.-Z., Wan, X.-F., Hua, Y., Li, L., Zhou, J., Wang, X., Hou, F., Chen, J., Zou, J. et al. (2020) Are pangolins the intermediate host of the 2019 novel coronavirus (SARS-CoV-2)? PloS Pathogens, 16, e1008421.

45. Yang, Z.H. and Rannala, B. (2012) Molecular phylogenetics: principles and practice. Nat. Rev. Genet., 13, 303–314.

46. Harrison, C.J. and Langdale, J.A. (2006) A step by step guide to phylogeny reconstruction. Plant J, 45, 561–572.

47. Hunt, T. and Vogler, A.P. (2008) A protocol for large-scale rRNA sequence analysis: Towards a detailed phylogeny of Coleoptera. Mol. Phylogenet. Evol., 47, 289–301.

48. Hall, B.G. (2013) Building phylogenetic trees from molecular data with MEGA. Mol. Biol. Evol., 30, 1229–1235.

49. Lemmon, E.M. and Lemmon, A.R. (2013) High-throughput genomic data in systematics and phylogenetics. Annu. Rev. Ecol. Evol. Syst., 44, 99–121.

50. O’Halloran, D. (2014) A oractical guide to phylogenetics for nonexperts. J. Vis. Exp., 14.

51. Wilding, M., Nachtschatt, M., Speight, R. and Scott, C. (2017) An improved and general streamlined phylogenetic protocol applied to the fatty acid desaturase family. Mol. Phylogenet. Evol., 115, 50–57.

52. Morrison, D.A. (2015) Is sequence alignment an art or a science? Syst. Bot., 40, 14–26.

53. Jermiin, L.S., Lovell, D.R., Misof, B., Foster, P.G. and Robinson, J. (2020) Detecting heterogeneous evolutionary processes across aligned sequence data. Syst. Biol., **(in review)**, https://doi.org/10.1101/828996.

54. Castresana, J. (2000) Selection of conservative blocks from multiple alignments for their use in phylogenetic analysis. Mol. Biol. Evol., 17, 540–552.

55. Talavera, G. and Castresana, J. (2007) Improvement of phylogenies after removing divergent and ambiguously aligned blocks from protein sequence alignments. Syst. Biol., 56, 564–577.

56. Dress, A.W.M., Flamm, C., Fritzsch, G., Grunewald, S., Kruspe, M., Prohaska, S.J. and Stadler, P.F. (2008) Noisy: identification of problematic columns in multiple sequence alignments. Algorithms for Molecular Biology, 3, 7.

57. Hartmann, S. and Vision, T.J. (2008) Using ESTs for phylogenomics: can one accurately infer a phylogenetic tree from a gappy alignment? BMC Evol. Biol., 8, 95.

58. Misof, B. and Misof, K. (2009) A Monte Carlo approach successfully identifies randomness in multiple sequence alignments: a more objective means of data exclusion. Syst. Biol., 58, 21–34.

59. Capella-Gutierrez, S., Silla-Martinez, J.M. and Gabaldon, T. (2009) trimAl: a tool for automated alignment trimming in large-scale phylogenetic analyses. Bioinformatics, 25, 1972–1973.

60. Kück, P., Meusemann, K., Dambach, J., Thormann, B., von Reumont, B.M., Wägele, J.W. and Misof, B. (2010) Parametric and non-parametric masking of randomness in sequence alignments can be improved and leads to better resolved trees. Front. Zool., 7, 10.

61. Criscuolo, A. and Gribaldo, S. (2010) BMGE (Block Mapping and Gathering with Entropy): a new software for selection of phylogenetic informative regions from multiple sequence alignments. BMC Evol. Biol., 10, 210.

62. Penn, O., Privman, E., Landan, G., Graur, D. and Pupko, T. (2010) An alignment confidence score capturing robustness to guide tree uncertainty. Mol. Biol. Evol., 27, 1759–1767.

63. Wu, M.T., Chatterji, S. and Eisen, J.A. (2012) Accounting for alignment uncertainty in phylogenomics. PLoS One, 7, e30288.

64. Wong, T.F.K., Kalyaanamoorthy, S., Meusemann, K., Yeates, D.K., Misof, B. and Jermiin, L.S. (2020) A minimum reporting standard for multiple sequence alignments. NAR Genom. Bioinf., 2, lqaa024.

65. Tan, G., Muffato, M., Ledergerber, C., Herrero, J., Goldman, N., Gil, M. and Dessimoz, C. (2015) Current methods for automated filtering of multiple sequence alignments frequently worsen single-gene phylogenetic inference. Syst. Biol., 64, 778–791.

66. Bryant, D., Galtier, N. and Poursat, M.-A. (2005) In Gascuel, O. (ed.), Mathematics of Evolution and Phylogeny. Oxford University Press, Oxford, pp. 33–62.

67. Jayaswal, V., Jermiin, L.S. and Robinson, J. (2005) Estimation of phylogeny using a general Markov model. Evol. Bioinform., 1, 62–80.

68. Ababneh, F., Jermiin, L.S. and Robinson, J. (2006) Generation of the exact distribution and simulation of matched nucleotide sequences on a phylogenetic tree. J. Math. Model. Algor., 5, 291–308.

69. Swofford, D.L. (2003). 4 ed. Sinauer Associates, Sunderland, Massachusetts.

70. Paradis, E., Claude, J. and Strimmer, K. (2004) APE: Analyses of phylogenetics and evolution in R language. Bioinformatics, 20, 289–290.

71. Felsenstein, J. (2005). 3.6 ed. Distributed by the author, Seattle.

72. Popescu, A.A., Huber, K.T. and Paradis, E. (2012) ape 3.0: New tools for distancebased phylogenetics and evolutionary analysis in R. Bioinformatics, 28, 1536–1537.

73. Kumar, S., Stecher, G. and Tamura, K. (2016) MEGA7: Molecular evolutionary genetics analysis version 7.0 for bigger datasets. Mol. Biol. Evol., 33, 1870–1874.

74. Xia, X.H. (2017) DAMBE6: New tools for microbial genomics, phylogenetics, and molecular evolution. J. Hered., 108, 431–437.

75. Knight, R., Maxwell, P., Birmingham, A., Carnes, J., Caporaso, J.G., Easton, B.C., Eaton, M., Hamady, M., Lindsay, H., Liu, Z.Z. et al. (2007) PyCogent: a toolkit for making sense from sequence. Gen. Biol., 8, 16.

76. Guindon, S., Dufayard, J.F., Lefort, V., Anisimova, M., Hordijk, W. and Gascuel, O. (2010) New algorithms and methods to estimate maximum-likelihood phylogenies: Assessing the performance of PhyML 3.0. Syst. Biol., 59, 307–321.

77. Bazinet, A.L., Zwickl, D.J. and Cummings, M.P. (2014) A gateway for phylogenetic analysis powered by grid computing featuring GARLI 2.0. Syst. Biol., 63, 812–818.

78. Stamatakis, A. (2014) RAxML version 8: A tool for phylogenetic analysis and postanalysis of large phylogenies. Bioinformatics, 30, 1312–1313.

79. Nguyen, L.-T., Schmidt, H.A., Von Haeseler, A. and Minh, B.Q. (2015) IQ-TREE: A fast and effective stochastic algorithm for estimating maximum-likelihood phylogenies. Mol. Biol. Evol., 32, 268–274.

80. Lartillot, N., Lepage, T. and Blanquart, S. (2009) PhyloBayes 3: a Bayesian software package for phylogenetic reconstruction and molecular dating. Bioinformatics, 25, 2286–2288.

81. Ronquist, F., Teslenko, M., van der Mark, P., Ayres, D.L., Darling, A., Hohna, S., Larget, B., Liu, L., Suchard, M.A. and Huelsenbeck, J.P. (2012) MrBayes 3.2: Efficient Bayesian phylogenetic inference and model choice across a large model space. Syst. Biol., 61, 539–542.

82. Lartillot, N., Rodrigue, N., Stubbs, D. and Richer, J. (2013) PhyloBayes MPI: Phylogenetic Reconstruction with Infinite Mixtures of Profiles in a Parallel Environment. Syst. Biol., 62, 611–615.

83. Bouckaert, R., Heled, J., Kuhnert, D., Vaughan, T., Wu, C.H., Xie, D., Suchard, M.A., Rambaut, A. and Drummond, A.J. (2014) BEAST 2: a software platform for Bayesian evolutionary analysis. PLoS Comp. Biol., 10, 6.

84. Höhna, S., Landis, M.J., Heath, T.A., Boussau, B., Lartillot, N., Moore, B.R., Huelsenbeck, J.P. and Ronquist, F. (2016) RevBayes: Bayesian phylogenetic inference using graphical models and an interactive model-specification language. Syst. Biol., 65, 726–736.

85. Ogilvie, H.A., Heled, J., Xie, D. and Drummond, A.J. (2016) Computational performance and statistical accuracy of *BEAST and comparisons with other methods. Syst. Biol., 65, 381–396.

86. Suchard, M.A., Lemey, P., Baele, G., Ayres, D.L., Drummond, A.J. and Rambaut, A. (2018) Bayesian phylogenetic and phylodynamic data integration using BEAST 1.10. Virus Evol., 4, vey016.

87. Bouckaert, R., Vaughan, T.G., Barido-Sottani, J., Duchéne, S., Fourment, M., Gavryushkina, A., Heled, J., Jones, G., Kühnert, D., De Maio, N. et al. (2019) BEAST 2.5: An advanced software platform for Bayesian evolutionary analysis. PLoS Comp. Biol., 15, e1006650.

88. Goloboff, P.A., Farris, J.S. and Nixon, K.C. (2008) TNT, a free program for phylogenetic analysis. Cladistics, 24, 774–786.

89. Goloboff, P.A. and Catalano, S.A. (2016) TNT version 1.5, including a full implementation of phylogenetic morphometrics. Cladistics, 32, 221–238.

90. White, W.T.J. and Holland, B.R. (2011) Faster exact maximum parsimony search with XMP. Bioinformatics, 27, 1359–1367.

91. Posada, D. and Crandall, K.A. (1998) MODELTEST: testing the model of DNA substitution. Bioinformatics, 14, 817–818.

92. Chiotis, M., Jermiin, L.S. and Crozier, R.H. (2000) A molecular framework for the phylogeny of the ant subfamily Dolichoderinae. Mol. Phylogenet. Evol., 17, 108–116.

93. Abascal, F., Zardoya, R. and Posada, D. (2005) ProtTest: selection of best-fit models of protein evolution. Bioinformatics, 21, 2104–2105.

94. Keane, T.M., Creevey, C.J., Pentony, M.M., Naughton, T.J. and McInerney, J.O. (2006) Assessment of methods for amino acid matrix selection and their use on empirical data shows that ad hoc assumptions for choice of matrix are not justified. BMC Evol. Biol., 6, 29.

95. Posada, D. (2006) ModelTest Server: a web-based tool for the statistical selection of models of nucleotide substitution online. Nucl. Acid. Res., 34, W700–W703.

96. Posada, D. (2008) jModelTest: Phylogenetic model averaging. Mol. Biol. Evol., 25, 1253–1256.

97. Darriba, D., Taboada, G.L., Doallo, R. and Posada, D. (2011) ProtTest 3: fast selection of best-fit models of protein evolution. Bioinformatics, 27, 1164–1165.

98. Darriba, D., Taboada, G.L., Doallo, R. and Posada, D. (2012) jModelTest 2: More models, new heuristics and parallel computing. Nature Meth., 9, 772.

99. Lanfear, R., Calcott, B., Ho, S.Y.W. and Guindon, S. (2012) Partitionfinder: combined selection of partitioning schemes and substitution models for phylogenetic analyses. Mol. Biol. Evol., 29, 1695–1701.

100. Santorum, J.M., Darriba, D., Taboada, G.L. and Posada, D. (2014) jmodeltest.org: selection of nucleotide substitution models on the cloud. Bioinformatics, 30, 1310–1311.

101. Whelan, S., Allen, J.E., Blackburne, B.P. and Talavera, D. (2015) ModelOMatic: Fast and automated model selection between RY, nucleotide, amino acid, and codon substitution models. Syst. Biol., 64, 42–55.

102. Lefort, V., Longueville, J.E. and Gascuel, O. (2017) SMS: Smart model selection in PhyML. Mol. Biol. Evol., 34, 2422–2424.

103. Minh, B.Q., Nguyen, M.A.T. and von Haeseler, A. (2013) Ultrafast approximation for phylogenetic bootstrap. Mol. Biol. Evol., 30, 1188–1195.

104. Larget, B. and Simon, D. (1999) Markov chain Monte Carlo algorithms for the Bayesian analysis of phylogenetic trees. Mol. Biol. Evol., 16, 750–759.

105. Goremykin, V.V., Nikiforova, S.V., Biggs, P.J., Zhong, B.J., Delange, P., Martin, W., Woetzel, S., Atherton, R.A., McLenachan, P.A. and Lockhart, P.J. (2013) The evolutionary root of flowering plants. Syst. Biol., 62, 50–61.

106. Drew, B.T., Ruhfel, B.R., Smith, S.A., Moore, M.J., Briggs, B.G., Gitzendanner, M.A., Soltis, P.S. and Soltis, D.E. (2014) Another look at the root of the angiosperms reveals a familiar tale. Syst. Biol., 63, 368–382.

107. Goremykin, V.V., Nikiforova, S.V., Cavalieri, D., Pindo, M. and Lockhart, P. (2015) The root of flowering plants and total evidence. Syst. Biol., 64, 879–891.

108. Rokas, A., Krüger, D. and Carroll, S.B. (2005) Animal evolution and the molecular signature of radiations compressed in time. Science, 310, 1933–1938.

109. Catullo, R.A. and Oakeshott, J.G. (2014) Problems with data quality in the reconstruction of evolutionary relationships in the Drosophila melanogaster species group: Comments on Yang et al. (2012). Mol. Phylogenet. Evol., 78, 275–276.

110. Ashkenazy, H., Sela, I., Karin, E.L., Landan, G. and Pupko, T. (2019) Multiple sequence alignment averaging Improves phylogeny reconstruction. Syst. Biol., 68, 117–130.

111. Morrison, D.A. (2006) Multiple sequence alignment for phylogenetic purposes. Aust. Syst. Bot., 19, 479–539.

112. Golubchik, T., Wise, M.J., Easteal, S. and Jermiin, L.S. (2007) Mind the gaps: Evidence of bias in estimates of multiple sequence alignments. Mol. Biol. Evol., 24, 2433–2442.

113. Morrison, D.A. (2009) A framework for phylogenetic sequence alignment. Plant Syst. Evol., 282, 127–149.

114. Morrison, D.A. (2009) Why would phylogeneticists ignore computerized sequence alignment? Syst. Biol., 58, 150–158.

115. Sievers, F., Wilm, A., Dineen, D., Gibson, T.J., Karplus, K., Li, W.Z., Lopez, R., McWilliam, H., Remmert, M., Soding, J. et al. (2011) Fast, scalable generation of high-quality protein multiple sequence alignments using Clustal Omega. Mol. Syst. Biol., 7, 539.

116. Thompson, J.D., Linard, B., Lecompte, O. and Poch, O. (2011) A comprehensive benchmark study of multiple sequence alignment methods: Current challenges and future perspectives. PloS One, 6, 14.

117. Chatzou, M., Magis, C., Chang, J.M., Kemena, C., Bussotti, G., Erb, I. and Notredame, C. (2016) Multiple sequence alignment modeling: methods and applications. Brief. Bioinf., 17, 1009–1023.

118. Chowdhury, B. and Garai, G. (2017) A review on multiple sequence alignment from the perspective of genetic algorithm. Genomics, 109, 419–431.

119. Jordan, G. and Goldman, N. (2011) The effects of alignment error and alignment filtering on the sitewise detection of positive selection. Mol. Biol. Evol., 29, 1125–1139.

120. Vialle, R.A., Tamuri, A.U. and Goldman, N. (2018) Alignment modulates ancestral sequence reconstruction accuracy. Mol. Biol. Evol., 35, 1783–1797.

121. Blackburne, B.P. and Whelan, S. (2012) Class of multiple sequence alignment algorithm affects genomic analysis. Mol. Biol. Evol., 30, 642–653.

122. Ho, S.Y.W. and Jermiin, L.S. (2004) Tracing the decay of the historical signal in biological sequence data. Syst. Biol., 53, 623–637.

123. Jermiin, L.S., Ho, S.Y.W., Ababneh, F., Robinson, J. and Larkum, A.D.W. (2004) The biasing effect of compositional heterogeneity on phylogenetic estimates may be underestimated. Syst. Biol., 53, 638–643.

124. Cooper, E.D. (2014) Overly simplistic substitution models obscure green plant phylogeny. Trends Plant Sci., 19, 576–582.

125. Jermiin, L.S., Poladian, L. and Charleston, M.A. (2005) Evolution - Is the “Big Bang” in animal evolution real? Science, 310, 1910–1911.

126. Winking, J. (2018) Exploring the great schism in the Social Sciences: Confirmation bias and the interpretation of results relating to biological influences on human behavior and psychology. Evol. Psychol., 16, 10.

127. Tuller, T. and Mossel, E. (2011) Co-evolution is incompatible with the Markov assumption in phylogenetics. IEEE/ACM Transactions on Computational Biology and Bioinformatics, 8, 1667–1670.

128. Vera-Ruiz, V.A., Lau, K.W., Robinson, J. and Jermiin, L.S. (2014) Statistical tests to identify appropriate types of nucleotide sequence recoding in molecular phylogenetics. BMC Bioinformatics, 15 (Suppl. 2), S8.

129. Vera-Ruiz, V.A., Robinson, J. and Jermiin, L.S. (2020) A likelihood-ratio test for lumpability of phylogenetic data: Is the Markovian property of an evolutionary process retained in recoded DNA? (in preperation).

130. Nasrallah, C.A., Mathews, D.H. and Huelsenbeck, J.P. (2011) Quantifying the impact of dependent evolution among sites in phylogenetic inference. Syst. Biol., 60, 60–73.

131. Siepel, A. and Haussler, D. (2004) Phylogenetic estimation of context-dependent substitution rates by maximum likelihood. Mol. Biol. Evol., 21, 468–488.

132. Lindsay, H., Yap, V.B., Ying, H. and Huttley, G.A. (2008) Pitfalls of the most commonly used models of context dependent substitution. Biol. Direct, 3, 52.

133. Shapiro, B., Rambaut, A. and Drummond, A.J. (2005) Choosing appropriate substitution models for the phylogenetic analysis of protein-coding sequences. Mol. Biol. Evol., 23, 7–9.

134. Tavaré, S. (1986) Some probabilistic and statistical problems on the analysis of DNA sequences. Lect. Math. Life Sci., 17, 57–86.

135. Lanave, C. and Pesole, G. (1993) Stationary MARKOV processes in the evolution of biological macromolecules. Binary, 5, 191–195.

136. Rzhetsky, A. and Nei, M. (1995) Tests of applicability of several substitution models for DNA sequence data. Mol. Biol. Evol., 12, 131–151.

137. Weiss, G. and von Haeseler, A. (2003) Testing substitution models within a phylogenetic tree. Mol. Biol. Evol., 20, 572–578.

138. Ababneh, F., Jermiin, L.S., Ma, C. and Robinson, J. (2006) Matched-pairs tests of homogeneity with applications to homologous nucleotide sequences. Bioinformatics, 22, 1225–1231.

139. Ho, J.W.K., Adams, C.E., Lew, J.B., Matthews, T.J., Ng, C.C., Shahabi-Sirjani, A., Tan, L.H., Zhao, Y., Easteal, S., Wilson, S.R. et al. (2006) SeqVis: Visualization of compositional heterogeneity in large alignments of nucleotides. Bioinformatics, 22, 2162–2163.

140. Jermiin, L.S., Jayaswal, V., Ababneh, F. and Robinson, J. (2008) In Keith, J. (ed.), Bioinformatics, Volume 1: Data, sequence analysis, and evolution. Humana Press, Totowa, NJ, Vol. I, pp. 331–364.

141. Jermiin, L.S., Jayaswal, V., Ababneh, F.M. and Robinson, J. (2017) In Keith, J. (ed.), Bioinformatics: Volume 1: Data, Sequence Analysis, and Evolution. Humana Press, Totowa, NJ, pp. 379–420.

142. Naser-Khdour, S., Minh, B.Q., Zhang, W., A., S.E. and R., L. (2019) The prevalence and impact of model violations in phylogenetic analysis. Gen. Biol. Evol., 11, 3341–3352.

143. Kedzierska, A.M., Drton, M., Guigó, R. and Casanellas, M. (2011) SPIn: model selection for phylogenetic mixtures via linear invariants. Mol. Biol. Evol., 29, 929–937.

144. Lockhart, P.J. and Steel, M. (2005) A tale of two processes. Syst. Biol., 54, 948–955.

145. Eigen, M., Winkler-Oswatitsch, R. and Dress, A. (1988) Statistical geometry in sequence space: A method of quantitative comparative sequence analysis. Proc. Natl. Acad. Sci. USA, 85, 5913–5917.

146. Holland, B.R., Huber, K.T., Dress, A. and Moulton, V. (2002) δ plots: a tool for analyzing phylogenetic distance data. Mol. Biol. Evol., 19, 2051–2059.

147. Jermiin, L.S. and Misof, B. (2020) Measuring historical and compositional signals in phylogenetic data. In prep., https://doi.org/10.1101/2020.1101.1103.894097.

148. Townsend, J.P. (2007) Profiling phylogenetic informativeness. Syst. Biol., 56, 222–231.

149. López-Giráldez, F. and Townsend, J.P. (2011) PhyDesign: an online application for profiling phylogenetic informativeness. BMC Evol. Biol., 11, 152.

150. Dornburg, A., Fisk, N., Tamagnam, J. and Townsend, J.P. (2016) PhyInformR: phylogenetic experimental design and phylogenomic data exploration in R. BMC Evol. Biol., 16, 262.

151. Barry, D. and Hartigan, J.A. (1987) Statistical analysis of hominoid molecular evolution. Stat. Sci., 2, 191–210.

152. Reeves, J. (1992) Heterogeneity in the substitution process of amino acid sites of proteins coded for by the mitochondrial DNA. J. Mol. Evol., 35, 17–31.

153. Steel, M.A., Lockhart, P.J. and Penny, D. (1993) Confidence in evolutionary trees from biological sequence data. Nature, 364, 440–442.

154. Lake, J.A. (1994) Reconstructing evolutionary trees from DNA and protein sequences: paralinear distances. Proc. Natl. Acad. Sci. USA, 91, 1455–1459.

155. Lockhart, P.J., Steel, M.A., Hendy, M.D. and Penny, D. (1994) Recovering evolutionary trees under a more realistic model of sequence evolution. Mol. Biol. Evol., 11, 605–612.

156. Steel, M.A. (1994) Recovering a tree from the leaf colourations it generates under a Markov model. Appl. Math. Lett., 7, 19–23.

157. Galtier, N. and Gouy, M. (1995) Inferring phylogenies from DNA sequences of unequal base compositions. Proc. Natl. Acad. Sci. USA, 92, 11317–11321.

158. Steel, M.A., Lockhart, P.J. and Penny, D. (1995) A frequency-dependent significance test for parsimony. Mol. Phylogenet. Evol., 4, 64–71.

159. Yang, Z. and Roberts, D. (1995) On the use of nucleic acid sequences to infer early branches in the tree of life. Mol. Biol. Evol., 12, 451–458.

160. Gu, X. and Li, W.-H. (1996) Bias-corrected paralinear and logdet distances and tests of molecular clocks and phylogenies under nonstationary nucleotide frequencies. Mol. Biol. Evol., 13, 1375–1383.

161. Galtier, N. and Gouy, M. (1998) Inferring pattern and process: maximum-likelihood implementation of a nonhomogenous model of DNA sequence evolution for phylogenetic analysis. Mol. Biol. Evol., 15, 871–879.

162. Gu, X. and Li, W.-H. (1998) Estimation of evolutionary distances under stationary and nonstationary models of nucleotide substitution. Proc. Natl. Acad. Sci. USA, 95, 5899–5905.

163. Galtier, N., Tourasse, N. and Gouy, M. (1999) A nonhyperthermophilic common ancestor to extant life forms. Science, 283, 220–221.

164. Tamura, K. and Kumar, S. (2002) Evolutionary distance estimation under heterogeneous substitution pattern among lineages. Mol. Biol. Evol., 19, 1727–1736.

165. Foster, P.G. (2004) Modelling compositional heterogeneity. Syst. Biol., 53, 485–495.

166. Thollesson, M. (2004) LDDist: a Perl module for calculating LogDet pair-wise distances for protein and nucleotide sequences. Bioinformatics, 20, 416–418.

167. Blanquart, S. and Lartillot, N. (2006) A Bayesian compound stochastic process for modeling nonstationary and nonhomogeneous sequence evolution. Mol. Biol. Evol., 23, 2058–2071.

168. Jayaswal, V., Robinson, J. and Jermiin, L.S. (2007) Estimation of phylogeny and invariant sites under the General Markov model of nucleotide sequence evolution. Syst. Biol., 56, 155–162.

169. Blanquart, S. and Lartillot, N. (2008) A site- and time-heterogeneous model of amino acid replacement. Mol. Biol. Evol., 25, 842–858.

170. Dutheil, J. and Boussau, B. (2008) Non-homogeneous models of sequence evolution in the Bio++ suite of libraries and programs. BMC Evol. Biol., 8, 255.

171. Jayaswal, V., Jermiin, L.S., Poladian, L. and Robinson, J. (2011) Two stationary, non-homogeneous Markov models of nucleotide sequence evolution. Syst. Biol., 60, 74–86.

172. Jayaswal, V., Ababneh, F., Jermiin, L.S. and Robinson, J. (2011) Reducing model complexity when the evolutionary process over an edge is modeled as a homogeneous Markov process. Mol. Biol. Evol., 28, 3045–3059.

173. Dutheil, J.Y., Galtier, N., Romiguier, J., Douzery, E.J.P., Ranwez, V. and Boussau, B. (2012) Efficient selection of branch-specific models of sequence evolution. Mol. Biol. Evol., 29, 1861–1874.

174. Zou, L.W., Susko, E., Field, C. and Roger, A.J. (2012) Fitting nonstationary generaltime-reversible models to obtain edge-lengths and frequencies for the Barry-Hartigan model. Syst. Biol., 61, 927–940.

175. Groussin, M., Boussau, B. and Gouy, M. (2013) A branch-heterogeneous model of protein evolution for efficient inference of ancestral sequences. Syst. Biol., 62, 523–538.

176. Holland, B.R., Jarvis, P.D. and Sumner, J.G. (2013) Low-parameter phylogenetic inference under the General Markov Model. Syst. Biol., 62, 78–92.

177. Goldman, N. (1993) Statistical tests of models of DNA substitution. J. Mol. Evol., 36, 182–198.

178. Rambaut, A. and Grassly, N.C. (1997) Seq-Gen: an application for the Monte Carlo simulation of DNA sequence evolution along phylogenetic trees. Comput. Appl. Biosci., 13, 235–238.

179. Fletcher, W. and Yang, Z. (2009) INDELible: a flexible simulator of biological sequence evolution. Mol. Biol. Evol., 26, 1879–1888.

180. Mallo, D., de Oliveira Martins, L. and Posada, D. (2016) SimPhy: phylogenomic simulation of gene, locus, and species trees. Syst. Biol., 65, 334–344.

181. Wang, H.C., Spencer, M., Susko, E. and Roger, A.J. (2007) Testing for covarion-like evolution in protein sequences. Mol. Biol. Evol., 24, 294–305.

182. Bollback, J.P. (2002) Bayesian model adequacy and choice in phylogenetics. Mol. Biol. Evol., 19, 1171–1180.

183. Susko, E. and Roger, A.J. (2020) On the use of information criteria for model selection in phylogenetics. Mol. Biol. Evol., 37, 549–562.

184. Dziak, J.J., Coffman, D.L., Lanza, S.T., Li, R. and Jermiin, L.S. (2020) Sensitivity and specificity of information criteria. Brief. Bioinf., 21, 533–565.

185. Duchêne, D.A., Duchêne, S. and Ho, S.Y.W. (2018) PhyloMAd: efficient assessment of phylogenomic model adequacy. Bioinformatics, 34, 2300–2301.

186. Kumar, S., Filipski, A.J., Battistuzzi, F.U., Pond, S.L.K. and Tamura, K. (2012) Statistics and truth in phylogenomics. Mol. Biol. Evol., 29, 457–472.

187. Holland, B.R. (2013) The rise of statistical phylogenetics. Aust. N. Zea. J. Stat., 55, 205–220.

188. Yang, Z. (1996) Among-site rate variation and its impact on phylogenetic analysis. Trends Ecol. Evol., 11, 367–372.

189. Yang, Z. (1995) A space-time process model for the evolution of DNA sequences. Genetics, 139, 993–1005.

190. Kubatko, L.S., Carstens, B.C. and Knowles, L.L. (2009) STEM: species tree estimation using maximum likelihood for gene trees under coalescence. Bioinformatics, 25, 971–973.

191. Heled, J. and Drummond, A.J. (2010) Bayesian inference of species trees from multilocus data. Mol. Biol. Evol., 27, 570–580.

192. Bryant, D., Bouckaert, R., Felsenstein, J., Rosenberg, N.A. and RoyChoudhury, A. (2012) Inferring species trees directly from biallelic genetic markers: Bypassing gene trees in a full coalescent analysis. Mol. Biol. Evol., 29, 1917–1932.

193. Weinreich, D.M., Delaney, N.F., DePristo, M.A. and Hartl, D.L. (2006) Darwinian evolution can follow only very few mutational paths to fitter proteins. Science, 312, 111–114.

194. Bridgham, J.T., Ortlund, E.A. and Thornton, J.W. (2009) An epistatic ratchet constrains the direction of glucocorticoid receptor evolution. Nature, 461, 515–519.

195. Noor, S., Taylor, M.C., Russell, R.J., Jermiin, L.S., Jackson, C.J., Oakeshott, J.G. and Scott, C. (2012) Intramolecular epistasis and the evolution of a new enzymatic function. PLoS One, 7, e39822.

196. Galtier, N. (2001) Maximum-likelihood phylogenetic analysis under a covarion-like model. Mol. Biol. Evol., 18, 866–873.

197. Huelsenbeck, J.P. (2002) Testing a covariotide model of DNA substitution. Mol. Biol. Evol., 19, 98–707.

198. Crotty, S.M., Minh, B.Q., Bean, N.G., Holland, B.R., Tuke, J., Jermiin, L.S. and von Haeseler, A. (2020) GHOST: Recovering historical signal from heterotachously evolved sequence alignments. Syst. Biol., 69, 249–264.

199. Jermiin, L.S., Ho, J.W.K., Lau, K.W. and Jayaswal, V. (2009) In Posada, D. (ed.), Bioinformatics for DNA Sequence Analysis. Humana Press, Totowa, NJ, pp. 65–91.

200. Kelly, C. (1994) A test of the Markovian model of DNA evolution. Biometrics, 50, 653–664.

201. Squartini, F. and Arndt, P.F. (2008) Quantifying the stationarity and time reversibility of the nucleotide substitution process. Mol. Biol. Evol., 25, 2525–2535.

202. Huson, D.H. and Bryant, D. (2006) Application of phylogenetic networks in evolutionary studies. Mol. Biol. Evol., 23, 254–267.

203. Bapteste, E., van Iersel, L., Janke, A., Kelchner, S., Kelk, S., McInerney, J.O., Morrison, D.A., Nakhleh, L., Steel, M., Stougie, L. et al. (2013) Networks: expanding evolutionary thinking. Trends Genet., 29, 439–441.

204. Xia, X.F., Xie, Z., Salemi, M., Chen, L. and Wang, Y. (2003) An index of substitution saturation and its application. Mol. Phylogenet. Evol., 26, 1–7.

205. Fischer, M. and Steel, M. (2009) Sequence length bounds for resolving a deep phylogenetic divergence. J. Theor. Biol., 256, 247–252.

206. Mossel, E., Steel, M. and Gascuel, O. (2005), Mathematics of Evolution and Phylogeny. Oxford University Press, New York, pp. 384–422.

